# Small in size but huge as reservoir – insights into the virome of European white-toothed shrews

**DOI:** 10.1101/2023.11.14.567014

**Authors:** Viola C. Haring, Benedikt Litz, Jens Jacob, Michael Brecht, Markus Bauswein, Julia Sehl-Ewert, Marta Heroldova, Donata Hoffmann, Rainer G. Ulrich, Martin Beer, Florian Pfaff

## Abstract

While the virome and immune system of bats and rodents have been extensively studied, comprehensive data are lacking for insectivores. Anthropogenic land use and outdoor recreational activities may lead to an expansion of the human-shrew interface with the risk of zoonotic infections, as reported for Borna disease virus 1. We investigated the virosphere of four white-toothed shrew species from Central Europe, addressing the One Health concept of spillover prevention. A high diversity of viruses was identified, including several co-infections. Whole genomes were generated for novel species of paramyxoviruses (n=3), nairoviruses (n=2) and hepevirus. Phylogenetically, they are closely related to WHO priority diseases, such as henipaviruses. High viral loads of orthoparamyxoviruses were detected in kidneys, in well-perfused organs for orthonairoviruses, and an association with liver and intestine was identified for orthohepevirus. Our study highlights the virus diversity present in shrews, not only in biodiversity hotspots but also in industrialised countries.

## INTRODUCTION

Knowledge of pathogen diversity in wildlife species is essential to be prepared for the next pandemic, a key task in modern virology.^1^ Current estimates suggest that 75% of emerging human pathogens originated from (wild) animals.^2,3^ Small mammals, especially rodents and bats, are known reservoirs of zoonotic viruses^4–6^, but little is known about the virosphere of insectivore species, especially shrews.^7^ Shrews (Mammalia: Eulipotyphla: Soricidae) are species-rich and phylogenetically ancient (>45 million years).^8^ Three subfamilies are defined within the family Soricidae: Soricinae (red-toothed shrews), Crocidurinae (white-toothed shrews), and Myosoricinae (African white-toothed shrews). At least 242 species of 10 genera with an almost global distribution belong to the Crocidurinae subfamily and the great diversity is increasing with the discovery of new species (**Figure 1**).^9^

**Figure 1:**
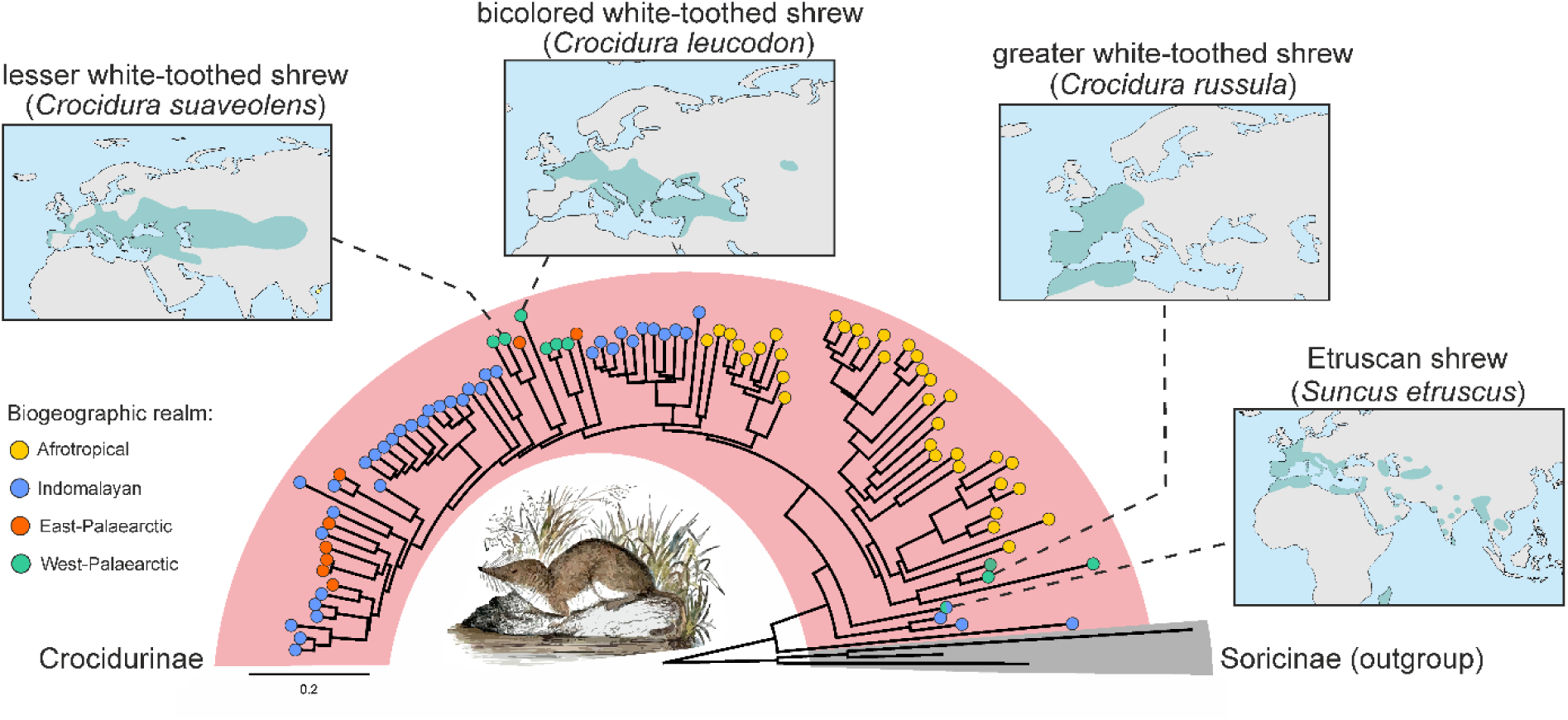
Phylogenetic relationships and biogeographic distribution of extant white-toothed shrews. The phylogenetic tree is based on all available *cytochrome b* sequences from white-toothed shrews (Crocidurinae) and a selected outgroup of red-toothed shrews (Soricinae) (IQ-TREE2; version 2.2.2.6). The biogeographic distribution of these animals can be broadly grouped into four realms: Afrotropical, Indomalayan, East- and West-Palearctic. The geographical range of the four Crocidurinae species that can be found in Europe, according to Wilson & Reeder 2017, is highlighted in the maps. Note the phylogenetic distances between *Suncus etruscus*, *Crocidura russula* and *Crocidura leucodon* / *Crocidura suaveolens*.

Primarily four synanthropic species of white-toothed shrews are found in Europe: bicolored white-toothed shrew (*Crocidura leucodon*), greater white-toothed shrew (*Crocidura russula*), lesser white-toothed shrew (*Crocidura suaveolens*), and Etruscan shrew (*Suncus etruscus*). ^8^ *Crocidura russula* originates from North Africa and is currently distributed across western Europe towards Fennoscandia and the Czech Republic.^10,11^ *Crocidura leucodon* is found from northern France through southern Europe to the Caspian Sea. The Etruscan shrew, one of the smallest recent living mammals with a body weight <2 g, is found mainly in southern Europe with a scattered distribution (**Figure 1**).^8^ The phylogenetic relationships among shrew species remain incompletely understood, with several species complexes, including the *C. suaveolens* sf. species complex, which shows a wide but fragmented distribution from the Atlantic coast to China.^8^

Interestingly, the number of new orthonairoviruses detected in shrews increases since the first report of Thiafora virus (TFAV) isolated from a *Crocidura* shrew in Senegal in 1971.^12^ Erve virus (ERVEV), which is thought to cause thunderclap headache in humans, was identified in *C. russula* from France.^12–14^ More recently, Lamusara virus (LMSV) and Lamgora virus (LMGV) have been described in the Goliath shrew (*Crocidura goliath*) from Gabon^15^, and Cencurut virus (CENV) in the Asian house shrew (*Suncus murinus*) from Singapore^16^, all of which belong to the Thiafora virus genogroup, which is closely related to zoonotic Crimean-Congo haemorrhagic fever virus (CCHFV). CCHFV causes highly contagious Crimean-Congo haemorrhagic fever in humans with a case fatality rate up to 40%.^17^ It is transmitted by ticks (*Hyalomma* spp.) or by direct contact to viraemic humans and animals. A small mammal reservoir for CCHFV has been discussed, but not identified. Ticks are now considered to be both reservoir and amplifying hosts.^17^

Recently, the zoonotic Langya virus (LayV, family *Paramyxoviridae*) was isolated from febrile human patients and detected in Ussuri white-toothed shrews (*Crocidura lasiura*) and Shantung white-toothed shrews (*Crocidura shantungensis*) in China.^18^ Gamak virus (GamV) and Daeryong virus have been identified in *C. lasiura* and *C. shantungensis* in Asia, respectively.^19^ A recent study in Belgium identified Melian virus (MeliV) in African *Crocidura grandiceps* and Denwin virus (DewV) in European *C. russula*.^20^ These all are related orthoparamyxoviruses of the genus *Henipavirus*, which includes the highly contagious and lethal zoonotic Hendra virus (HeV) and Nipah virus (NiV) detected in fruit bats in Australia and South-East Asia, respectively.^21,22^

At present, knowledge of pathogens in European shrews, especially white-toothed shrews, is limited, apart from intensive studies of *C. leucodon*, the natural reservoir for the zoonotic Borna disease virus 1 (BoDV-1), which causes fatal encephalitis in both humans and domestic animals.^23,24^

We investigated the virome of four white-toothed shrew species present in Europe using a straightforward sample pooling approach followed by high-throughput RNA sequencing and specific RT-qPCR confirmation and determination of the viral tissue distribution to identify potential transmission routes. Our study is thus one of the first to record the virome of white-toothed shrews in Europe. The surprisingly high number of novel viruses suggests a previously underestimated reservoir function of shrews, which might be even greater than that of sympatric rodent and bats species as postulated by Chen et al., 2023^25^, not only in subtropical but also in temperate regions.

## MATERIAL AND METHODS

### Sample selection and RNA extraction

A total of 19 bicolored white-toothed shrews (*Crocidura leucodon*), 16 greater white-toothed shrews (*Crocidura russula*), 6 lesser white-toothed shrews (*Crocidura suaveolens*) covering the known distribution of these species in Germany, captured between 2002 and 2021, and two additional *C. leucodon* collected in the Czech Republic in 2007 were selected. In addition, two Etruscan shrews (*Suncus etruscus*) from a German breeding colony were included in this study (Supplemental Figure S1 and Table S1). Identification of the shrew species was based on molecular analysis of the *cytochrome b* gene as described previously.^26^

First, organs were pooled per individual, consisting of small pieces of brain, lung, spleen, liver and kidney tissue, as available. These tissue pools were directly immersed in 1 ml QIAzol (Qiagen, Germany) and stored at -80°C until further processing. In addition, intestine tissue samples containing ingesta from several individuals of the same species were pooled and processed according to the individual tissue pools (Supplemental Table S1).

Tissue pools were homogenised for 2 min at 30 Hz using 5 mm steel beads on a TissueLyser II instrument (Qiagen, Germany). A volume of 0.2× chloroform (Carl Roth, Germany) was added to each reaction, mixed vigorously and centrifuged at 13,000×g for 10 min. The upper aqueous phase was further processed for total RNA extraction using the Agencourt RNAdvance Tissue Kit (Beckman Coulter, Germany) on a KingFisher Flex Purification System (Thermo Fisher Scientific, Germany) according to the manufacturer’s instructions.

### RNA library preparation and high throughput-sequencing

Total RNA quantity was measured using a Nanodrop ND1000 UV spectrophotometer (Peqlab, Germany) and total RNA quality was assessed using a 4150 TapeStation system (Agilent, Germany). In an attempt to reduce the amount of host-derived ribosomal RNA (rRNA), total RNA was treated with the “pan mammalia” riboPOOL ribosomal depletion kit (siTOOLs Biotech, Germany) according to the manufacturer’s instructions. The rRNA-depleted total RNA was then used for library preparation using the Collibri Stranded RNA Library Prep Kit for Illumina Systems (Invitrogen, Germany) according to the manufacturer’s instructions. Final libraries were quantified using a Qubit 2.0 fluorometer in conjunction with the Qubit dsDNA HS Assay-Kit (Invitrogen, Germany). The libraries were then pooled, submitted to CeGaT GmbH (Germany) and sequenced on a NovaSeq 6000 system (Illumina, USA) in 1×100 base pair (bp) mode.

### Sequence Data analysis

Raw reads were first trimmed for adapter contamination and poor quality using Trim Galore (version 0.6.10) in automatic adapter detection mode. Subsequently, host-specific background was then removed from the trimmed libraries using BBMap (version 39.01, k=13; ^27^) together with the combined genomic assemblies of *Crocidura indochinensis* (Indochinese white-toothed shrew, GCA_004027635.1), *Suncus etruscus* (GCF_024139225.1), *Sorex fumeus* (smokey shrew, GCA_026122425.1), *Sorex araneus* (common shrew, GCF_000181275.1) and *Cryptotis parvus* (North American least shrew, GCA_021461705.1) as reference. In addition, rRNA derived reads were removed using SortMeRNA (version 4.3.6; ^28^) with all rRNA entries of the SILVA database (release 138.1; ^29^) belonging to the taxon “Vertebrata” as reference.

The trimmed and host sequence-depleted libraries were individually assembled *de novo* using rnaSPAdes (version 3.15.5; ^30^). The metatranscriptomic pipeline SqueezeMeta (version 1.6.2; ^31^) was also used for *de novo* assembly, taxonomic classification and quantification. Specifically, SqueezeMeta was run with the option “-contiglen 400” in “seqmerge” mode, which merges individual assemblies into a single combined assembly prior to further processing. The assembly was then trimmed with regard to poly(A) and poly(T) sequences at the end or start of the contigs, using cutadapt (version 4.0; ^32^). This step will prevent unspecific mapping to poly(A)-tails. The trimmed *de novo* assembled contigs were then used for a final run of SqueezeMeta using the “-extassembly” option.

### Selection of complete viral genomes

Contigs that were classified as viral sequences and likely represented full genomes were selected based on their size from the SqueezeMeta assembly and compared with the rnaSPAdes assembly. For final quality check, the raw reads were mapped to the likely full genomes using Geneious Prime (version 2021.0.1) generic mapper. Open reading frame (ORF) annotation was done in Geneious Prime using appropriate references and the “Find ORFs” function. For selected samples we re-sequenced libraries and analysed them together with sequences obtained from individual kidney samples in order to improve coverage of the identified genomes.

### Rapid amplification of cDNA ends of the 5′end of whole genomes

Copy DNA was generated from total RNA using SuperScript III reverse transcriptase (Invitrogen, Germany) and 5′ Rapid Amplification of cDNA Ends (RACE) 2.0 system (Invitrogen, Germany) using a custom protocol to sequence the 5′ end of selected whole viral genomes was performed.

### Phylogenetic analysis of complete viral genomes

Viral sequences were aligned with publicly available reference sequences using MUSCLE (version 3.8.425). Maximum-likelihood phylogenetic trees were calculated using IQ-TREE2 (version 2.2.2.6; ^33^) with an automated model selection and each 100.000 ultra-fast bootstrap ^34^ and SH-aLRT ^35^ replicates.

In detail, for hepevirus phylogeny, we selected 36 representative genomes of the subfamily *Orthohepevirinae* and five genomes of fish hepeviruses (subfamily *Parahepevirinae)* as references for phylogenetic analysis. The first 450 aa (amino acids) of the ORF1 (non-structural polyprotein) were aligned and used for phylogeny.

For paramyxovirus phylogeny, we selected 54 representative genomes of the subfamily *Orthoparamyxovirinae* and one genome of the subfamily *Metaparamyxovirinae* as references for phylogenetic analysis. The amino acid sequences of the large protein (L, including RNA-directed RNA polymerase, capping and cap methylation activities) were aligned and used for phylogeny.

For nairovirus phylogeny, we selected 46 representative genomes of the genus *Orthonairovirus* and one genome of the genus *Shaspivirus* as references for phylogenetic analysis. The amino acid sequences of the large protein (L, large segment, containing an RNA-directed RNA polymerase domain) were aligned and used for phylogeny.

For bornavirus phylogeny, we selected 74 shrew and domestic animal derived genomes of the species *Orthobornavirus bornaense* (genus *Orthobornavirus*). Borna disease virus 2 (also species *Orthobornavirus bornaense*) was used as outgroup. Nucleotide sequences spanning the N, X and P protein were aligned and used for phylogenetic analysis.

### Virus-specific RT-qPCR

Primers and probes for RT-qPCR detection of viral RNA of the detected nairo-, paramyxo- and hepeviruses were designed using Primer3web (version 4.1.0; ^36^). The L ORF was targeted for nairoviruses and paramyxoviruses, and ORF3 for hepeviruses. For specific detection of BoDV-1, the BoDV-1-Mix1-FAM assay was used.^37^ A set of primers and probe targeting the ß-actin-2 gene was used as an internal control.^38^ Sequences are shown in Table S4. The RT-qPCR reactions were performed using the AgPath-ID One-Step RT-PCR Kit according to the manufacturer’s instructions and run on a CFX96 Touch Real-Time PCR Detection System (Bio- Rad, Germany) with the following protocol: 10 min at 45°C for reverse transcription, 10 min at 95°C for polymerase activation; 42 cycles of 15 s at 95°C, 20 s at 57°C (with fluorescence detection during this step), 30 s at 72°C.

### Tissue distribution of novel viruses

An organ panel was prepared from selected animals to assess the tissue distribution of viral RNA. Approximately 50 mg of tissue was homogenised in 500 µl phosphate-buffered saline (PBS) for 2 min at 30 Hz using 5 mm steel beads on a TissueLyser II instrument (Qiagen). Total nucleic acids were extracted using the Nucleo Mag Vet Kit (Macherey & Nagel, Germany) on a KingFisher Flex Purification System (Thermo Fisher Scientific) according to the manufacturer’s instructions.

### Virus isolation in cell culture

For cell culture isolation of Rasenna virus from *Suncus etruscus*, organ material was lysed in cell culture medium and used to inoculate Vero cells (CCLV-RIE 0228) or baby hamster kidney (BHK) 21 cells (CCLV-RIE 0179) in a TC12.5 format (serum-free cell culture medium plus antibiotics). The cell culture supernatant from each cell culture flask was used for passaging to achieve four consecutive passages. In addition, the cells were passaged again separately to obtain four consecutive passages. Organs used for the different isolation attempts included liver, spleen, heart, muscle, fat, skin, thoracic and cervical spinal cord.

## RESULTS AND DISCUSSION

### Overall virome analysis

The metagenomic analysis revealed the presence of a wide range of RNA viruses belonging to the orders *Bunyavirales*, *Mononegavirales*, *Hepelivirales*, *Picornavirales*, and *Stellavirales* (**Figure 2**). *Bunyavirales* and *Mononegavirales* were the most abundant orders in the individual-based organ pools, while *Picornavirales* and *Stellavirales* were predominantly detected in the species-based intestine pools. Organ pools provide the benefit of reduced sampling and sequencing bias due to non-homogeneous virus distribution in the different organs.

**Figure 2:**
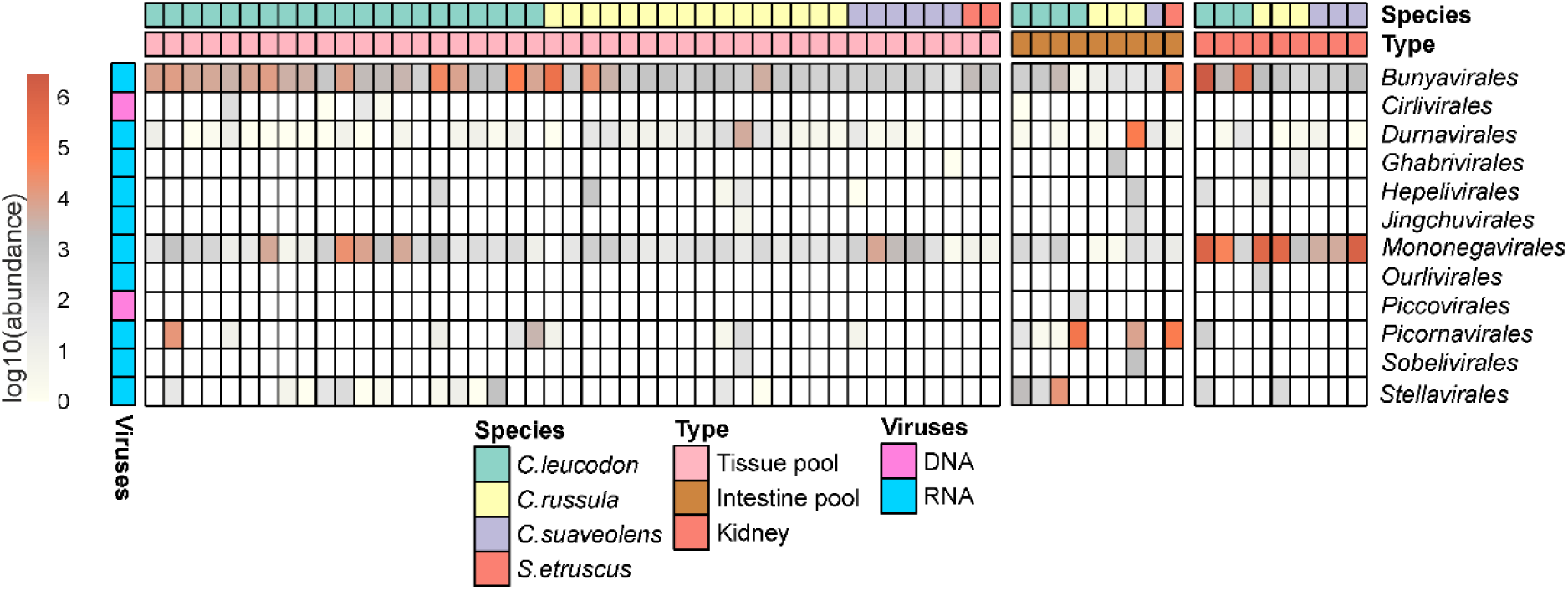
Viral diversity in different samples from white-toothed shrews. The heatmap shows the relative abundance of viral sequences sorted taxonomically by viral order. Note the abundances of the orders *Bunyavirales*, *Mononegavirales*, *Hepelivirales*, *Picornavirales*, and *Stellavirales*. *Bunyavirales* and *Mononegavirales* were the most abundant orders in the organ pools, while *Picornavirales* and *Stellavirales* were predominantly detected in the intestine pools. Based on the observed tissue distribution, kidney tissue from selected individuals was additionally sequenced and included in the analysis.

Subsequent analysis focused on virus genera with potentially zoonotic viruses with public health implications.^1,7^ In particular, we identified the whole genome sequences of novel paramyxo-, orthonairo- and orthohepeviruses, as well as several complete genome sequences of the zoonotic BoDV-1 and ERVEV.^14,23^ Virus-specific RT-qPCRs were designed in order to determine viral RNA tissue distribution. Based on the observed tissue distribution, kidney tissue from selected individuals was additionally sequenced and included in the analysis. The following sections summarise the results for each virus family.

### Detection and analysis of novel paramyxoviruses

Within the family *Paramyxoviridae* (order *Mononegavirales*) there are currently four subfamilies with 14 genera established.^20^ The subfamily *Orthoparamyxovirinae* comprises several viruses with a very high impact on human and animal health, such as members of the genera *Morbillivirus* (measles virus and rinderpest virus, the first successfully eradicated epizootic disease) and *Henipavirus* (NiV, HeV), with reoccurring outbreaks of NiV demonstrating dramatic case fatality rates of 40-70% including possible human-to-human transmission.^1,22,39^

Within the organ pools we identified genomes (**Figure 3A**) of diverse orthoparamyxoviruses that phylogenetically clustered within the genus *Henipavirus* (**Figure 3B**), forming a distinct shrew-dominated clade (**Figure 3B, C**).

**Figure 3:**
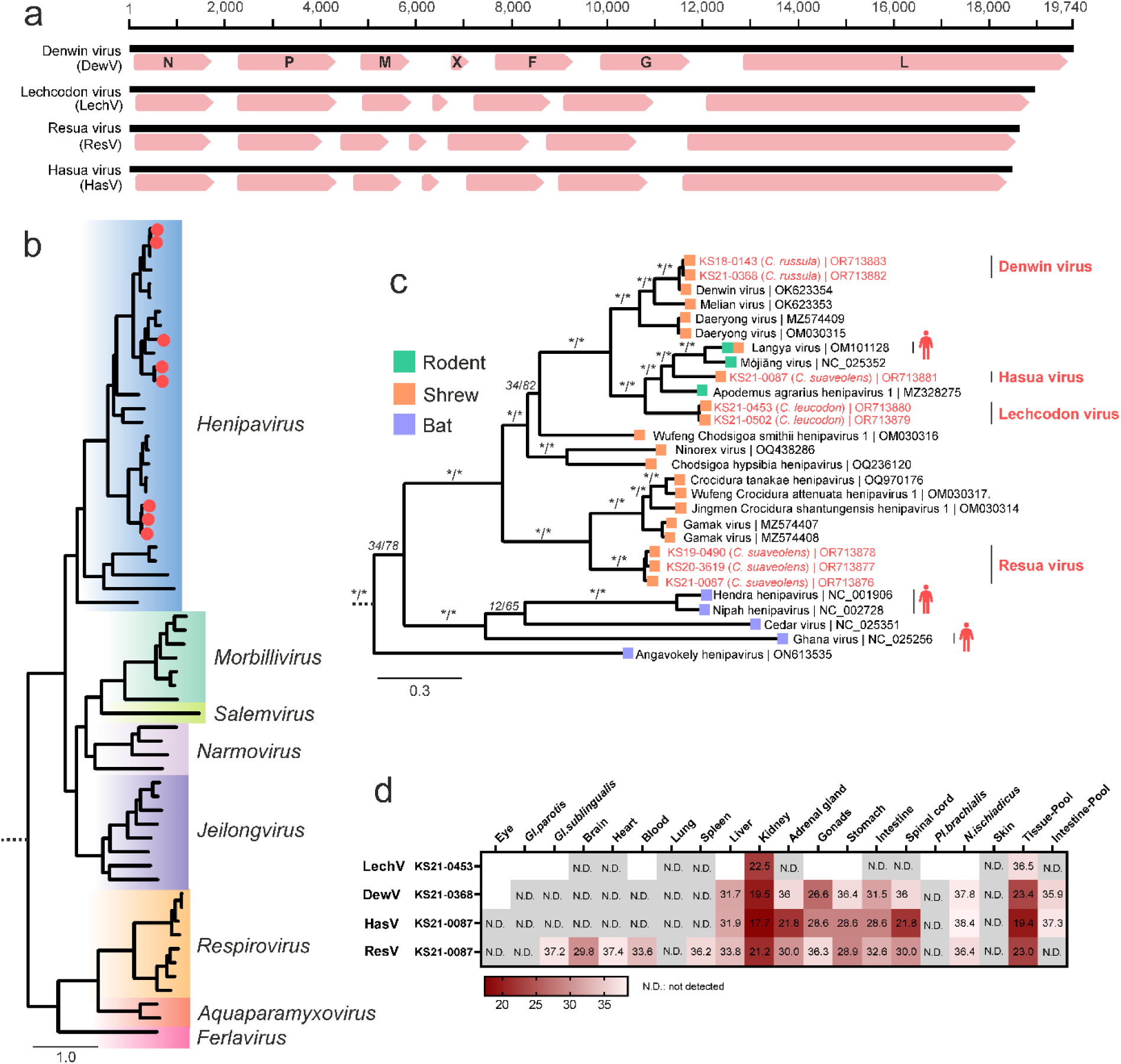
Detection and analysis of shrew-associated paramyxoviruses. **(a)** The genome structure of the novel paramyxoviruses was similar to that of Denwin virus, with the presence of the hypothetical open reading frame “X”, specific to shrew-derived paramyxoviruses.^20^ **(b)** For phylogenetic analysis, we selected 54 representative genomes from the *Orthoparamyxovirinae* subfamily, using *Metaparamyxovirinae* as outgroup. The amino acid sequences of the large protein (L, including RNA-directed RNA polymerase, capping and cap methylation activities) were aligned and used for phylogeny (IQ-TREE2; version 2.2.2.6). **(c)** Phylogenetic relationships within the genus *Henipavirus* with the novel whole genomes indicated in red, confirming the presence of a phylogenetically linked group of shrew-derived viruses that form a sister clade to the bat-borne henipaviruses. Host-association is indicated by colour of tips. Viruses with described zoonotic potential are highlighted with a human silhouette. Statistical support is shown for main branches using the format [SH-aLRT (%) / ultrafast bootstrap (%)]. Asterisks indicate statistical support ≥ 80% and ≥ 95% for ultrafast bootstrap and SH-aLRT, respectively. **(d)** Tissue distribution of paramyxovirus RNA using RT-qPCR specific for the L gene region. Results are given in cycle threshold (ct) values.

The novel Hasua virus (HasV) was identified in *C. suaveolens* (KS21-0087) from north-eastern Germany and was phylogenetically closely related to the zoonotic LayV and Mòjiāng virus (MojV) (compare Supplemental Table S2). Interestingly, we found sequences of another novel orthoparamyxovirus, Resua virus (ResV), in the same specimen (KS21-0087), suggesting co-infections. ResV was furthermore identified in two additional *C. suaveolens* from Germany (KS19-0490, KS20-3619). ResV phylogenetically clustered with a distant group of exclusively shrew-derived paramyxoviruses, such as GamV. Lechcodon virus (LechV) was detected in two *C. leucodon* from southern Germany (KS21-0502, KS21-0453) and grouped basal to HasV and LayV. Finally, sequences of the previously described DewV were detected in two *C. russula* (KS18-0143, KS21-0368), demonstrating its presence in Germany. In total, nine out of 16 *C. russula* were positive for DewV by RT-qPCR, indicating a wide geographical distribution of the virus (Supplemental Table S1).

Virus-specific RT-qPCR confirmed the presence of these viruses and viral RNA tropism was assessed, with high levels of viral RNA observed particularly in kidney tissue. Potential excretion and transmission via urine must be considered when establishing preventive measures (**Figure 3D**). Efficient transmission via urine was demonstrated for HeV and NiV, even allowing direct bat-to-human transmission for NiV through the consumption of urine-contaminated food.^22^ Otherwise, transmission of HeV and NiV from their fruit bat reservoir to humans requires an intermediate host, either horses or pigs, respectively.^22^

The zoonotic potential of these novel paramyxoviruses cannot be addressed in this study, as further *in vitro* and *in vivo* downstream characterisation are required.^6^ However, their striking phylogenetic proximity to known zoonotic agents (e. g. LayV, HeV, NiV) and to viruses that at least experimentally can infect human cells (GamV)^19^, clearly warrant such work. In any case, the findings indicate the need for biosafety considerations when handling these specimens. These newly identified paramyxoviruses confirm the presence of a phylogenetically related group of shrew-derived viruses that form a sister clade to the bat-borne henipaviruses and support the increasing number of globally distributed paramyxoviruses.^19,20,40^ This may ultimately lead to the establishment of a new shrew-borne genus within the *Orthoparamyxovirinae* subfamily.

### Detection and analysis of novel nairoviruses

The genus *Orthonairovirus* belongs to the family *Nairoviridae* of the order *Bunyavirales*. Orthonairoviruses are arthropod-borne, globally distributed viruses with a wide range of hosts, including mammals, birds, and even reptiles. In some cases, they can cause severe or even fatal disease in livestock and wildlife, with substantial economic and ecological implications.^3,16^ The reservoir species for many of these viruses have not been successfully identified yet and small mammals have been considered putative reservoirs or amplification hosts.^17^

In the sampled shrew organ pools, orthonairoviruses were highly abundant and detected in one quarter of the specimens across almost all species. Several phylogenetically distinct complete genomes could be deduced (**Figure 2, 4A**). These sequences were phylogenetically grouped within the Thiafora genogroup, a sister group to the CCHFV group, which includes the shrew-borne ERVEV, TFAV and CENV (**Figure 4B**).

**Figure 4:**
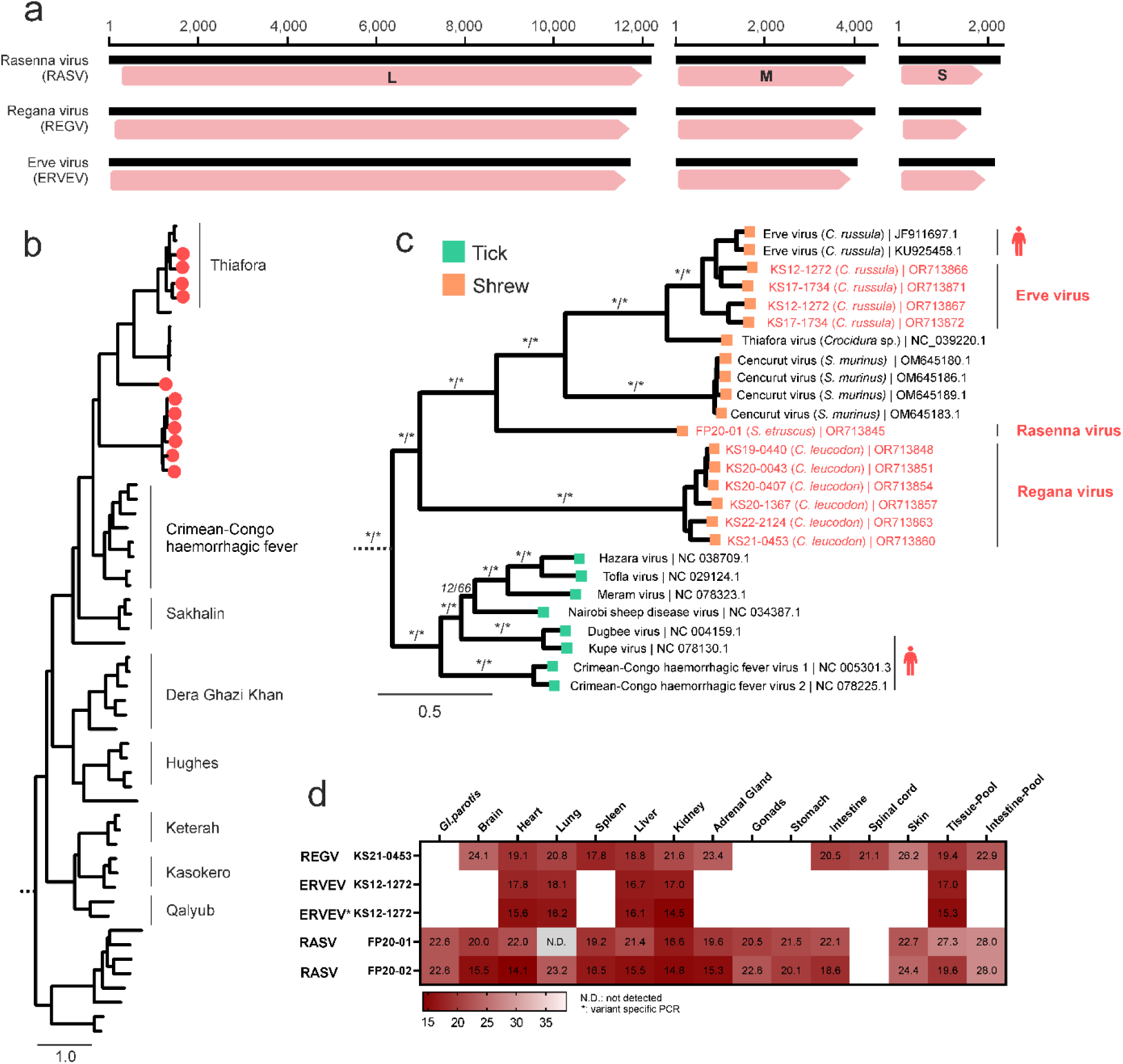
Detection and analysis of shrew-associated orthonairoviruses. **(a)** The segmented genome of the novel orthonairoviruses matched the size and number of the genomes of other members of the family *Nairovirida*e: the small (S) segment encoding for the nucleoprotein (N) and the non-structural NSs, the medium (M) segment encoding for glycoproteins and the large (L) segment encoding for the RNA-dependent RNA polymerase. **(b)** For the phylogeny of orthonairoviruses, we selected 46 representative genomes of the genus *Orthonairovirus* and *Shaspivirus* as outgroup. The amino acid sequences of the large protein (L) were aligned and used for phylogeny (IQ-TREE2; version 2.2.2.6). Novel genomes are indicated as red dots. **(c)** Detailed view of the phylogenetic relationships within the Crimean-Congo haemorrhagic fever and Thiafora genogroups. Newly generated whole genomes of Erve virus, Rasenna virus and Regana virus are shown in red. Host-association is indicated by colour. Viruses with described zoonotic potential are highlighted with a human silhouette. Statistical support is shown for main branches using the format [SH-aLRT (%) / ultrafast bootstrap (%)]. Asterisks indicate statistical support ≥ 80% and ≥ 95% for ultrafast bootstrap and SH-aLRT, respectively. **(d)** Viral RNA tissue distribution as determined by virus-specific RT-qPCRs. KS12-1272 was tested with two different primers and probe sets to differentiate between the two strains of Erve virus. Results are given in cycle threshold (ct) values.

In detail, whole genomes of the novel Regana virus (REGV) were identified in five *C. leucodon* (KS19-0440, KS20-0043, KS20-0407, KS20-1367, KS21-0453) across Germany and in one *C. leucodon* (KS22-2124) from the Czech Republic, forming a monophyletic cluster basal to the known viruses within the Thiafora genogroup (**Figure 4C** and Supplemental Table S2). The novel Rasenna virus (RASV) was identified in captive *S. etruscus* (FP20-1), clustering between REGV and CENV. Furthermore, we identified ERVEV in *C. russula* (KS12-1272, KS17-1734).

Virus-specific RT-qPCR confirmed the presence of the new virus genomes in the organ pools and showed a broad tissue distribution with all organs showing relatively high viral loads, especially the well-perfused organs. Liver tissue yielded the lowest cycle threshold (ct) values in all individuals (**Figure 4D**). These findings are in accordance with a viraemic status during the pathogenesis of orthonairoviruses and may indicate its circulation in the bloodstream. The putative role of ticks in the transmission of the orthonairoviruses detected remains a question for further study. However, the presence of RASV in captive *S. etruscus* from a well-established breeding colony suggests arthropod-independent transmission, as these animals were kept in a controlled ectoparasite-free environment.^41^ Vertical and efficient direct shrew-to-shrew transmission via scratching and biting during territorial fights may be assumed for the stable viral persistence in the colony.^8^

The zoonotic potential of these novel viruses is currently unknown, however ERVEV has been associated with reports of thunderclap headache in humans.^14,42^ The presence of genetically diverse ERVEV and the identification of two new shrew-borne orthonairoviruses (REGV in *C. leucodon* and RASV in *S. etruscus*) demonstrate the high diversity of orthonairoviruses in white-toothed shrews and increases the spectrum of potentially zoonotic nairoviruses.

### Detection and analysis of a novel hepevirus

Orthohepeviruses infect a wide range of animals including humans, pigs, rabbits, rodents, carnivores, bats and birds, but with exception of zoonotic viruses, they are generally highly host specific. Human hepatitis E virus (HEV) is faecal-orally transmitted and is a major cause of acute self-limiting hepatitis, particularly in developing countries. If transmitted vertically, it can cause early termination of pregnancy. The increasing number of food-borne cases of HEV-infections in industrialised countries is also of concern.^43^ Rodent-borne hepatitis E virus, which was first detected in Norway rats in Germany^44^, namely rat hepatitis E virus (ratHEV), has also been identified as a zoonotic agent worldwide. Severe chronic hepatitis can be induced by both pathogens in immunocompromised patients.^43^

Two closely related whole genomes of a novel hepevirus of the subfamily *Orthohepevirinae* were identified in two different specimens of *C. russula* (KS12-1272, KS21-0273) captured in western and eastern Germany (Supplemental Table S2). They show a genome organisation most similar to viruses of the genus *Paslahepevirus*, with an overlapping region of ORF2/ORF3 and the absence of ORF4, an open reading frame identified in viruses of the species *Rocahepevirus ratti* (ratHEV) (**Figure 5A**).^44^ This similarity in genome organisation is reflected in the phylogenetic position of shrewHEV, which clusters with strains of the genus *Paslahepevirus* well separated from strains of the genus *Rocahepevirus* (**Figure 5B**).

**Figure 5:**
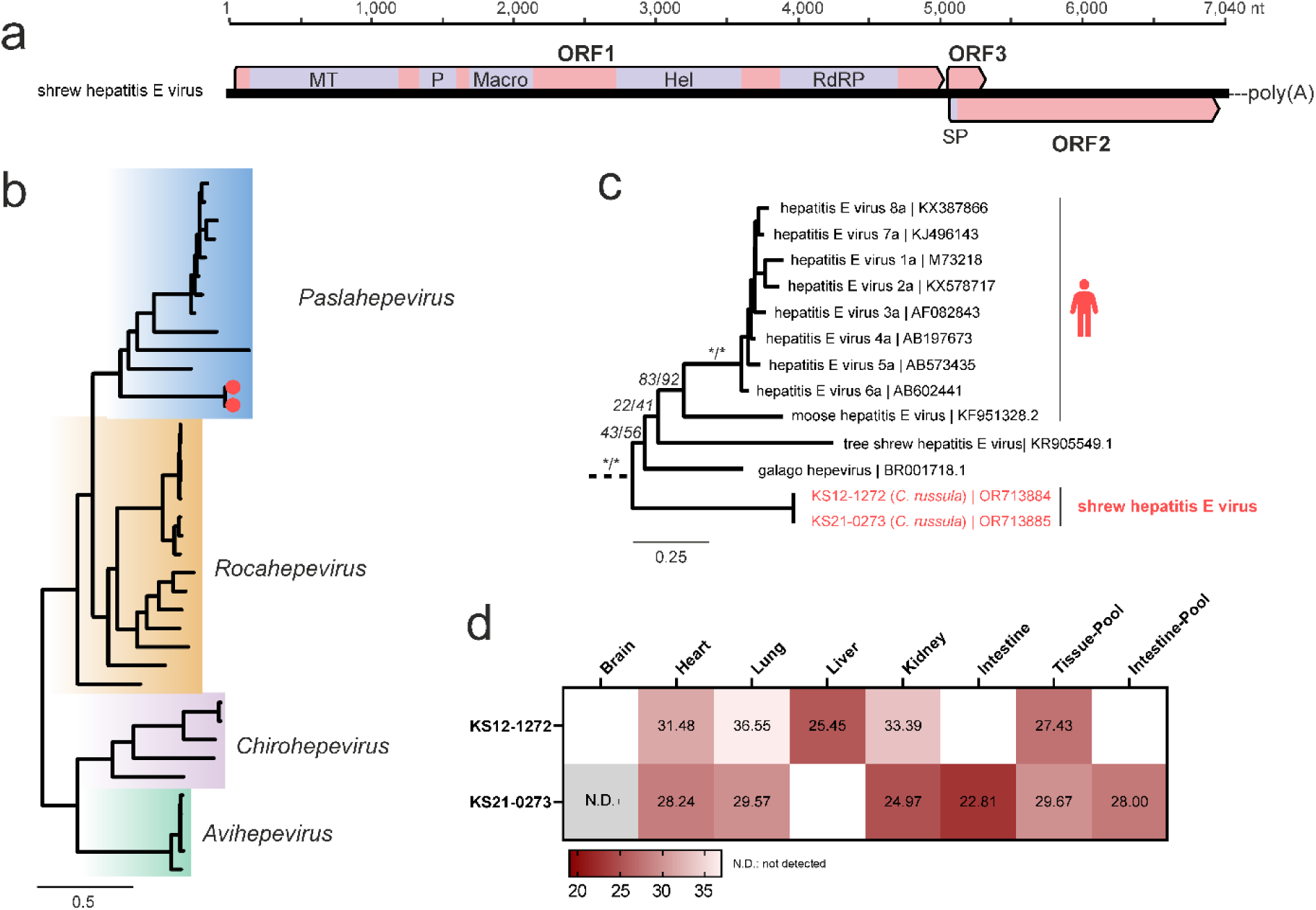
Detection and analysis of shrew-associated hepeviruses. **(a)** Genome structure of the novel shrew hepatitis E virus. **(b)** For the phylogenetic analysis of the novel hepevirus, 36 representative genomes of the subfamily *Orthohepevirinae* and five genomes of fish hepeviruses (subfamily *Parahepevirinae*) were selected as references. The first 450 aa of ORF1 (non-structural protein) were aligned and a phylogenetic tree was calculated (IQ-TREE2; version 2.2.2.6). Novel genomes are indicated as red dots. **(c)** Detailed view on the phylogenetic relations within the genus *Paslahepevirus*. Viruses with described zoonotic potential are highlighted with a human silhouette. The novel hepevirus sequences are indicated in red. Statistical support is shown for main branches using the format [SH-aLRT (%) / ultrafast bootstrap (%)]. Asterisks indicate statistical support ≥ 80% and ≥ 95% for ultrafast bootstrap and SH-aLRT, respectively. **(d)** Viral RNA tissue distribution of the novel shrewHEV in two *Crocidura russula* (KS12-1272, KS21-0273) as detected by virus-specific RT-qPCR. Results are given in cycle threshold (ct) values.

The highest viral RNA loads were detected in liver tissue of KS12-1272 and in kidney and intestine tissue of KS21-0273, suggesting a faecal-orally transmission, as described for HEV (**Figure 5C**). Due to its phylogenetic relationship with HEV, this commensal shrewHEV may also have zoonotic potential, pending confirmation in large-scale epidemiological studies.

### Detection and analysis of Borna disease virus 1

BoDV-1 belongs to the genus *Orthobornavirus* (family *Bornaviridae)*. It causes sporadic but highly lethal encephalitis in domestic animals, mainly horses, sheep and New World camelids, and has only been confirmed as zoonotic in 2018.^23,37^ The transmission from its reservoir, to dead-end hosts, its presence in the reservoir population, and the appearance of its endemic area are still poorly understood.

In this study, we generated seven new BoDV-1 complete genome sequences from four *C. leucodon* and, for the first time, from two *C. suaveolens* and one *C. russula* (Supplemental Figure S2 and Table S2). These new BoDV-1 sequences fall within the established phylogeographic clusters.^23^ The presence of BoDV-1 RNA in the tissue pools was confirmed by specific RT-qPCRs.

### Co-infection of different viruses

Several shrews in the study demonstrated co-infections with multiple viruses. For example, *C. russula* KS12-1272 was found to carry three viruses: the complete genome of shrewHEV, two different complete genomes of ERVEV, and it tested positive for DewV by RT-qPCR. Similarly, *C. suaveolens* KS21-0087 showed a triple infection, containing the complete genomes of two distinct paramyxoviruses (HasV and ResV) and BoDV-1. *Crocidura russula* KS21-0368 tested positive for both DewV and BoDV-1, while *C. suaveolens* KS20-3619 tested positive for BoDV-1 and ResV as detailed in Supplemental Figure S3.

In addition to co-infections with viruses from different taxonomic groups, sequencing revealed the co-occurrence of different variants of ERVEV in one specimen. This observation was subsequently confirmed using genome-specific RT-qPCR (**Figure 4D**). This finding suggests the potential for reassortment among these viruses, a process that can result in high genetic variability, especially in segmented viruses such as those of the order *Bunyavirales*.^17,45^

## CONCLUSION

Investigations of species-rich and phylogenetically ancient wildlife taxa such as shrews improves our understanding of global virus distribution.^46^ Revisions to existing taxonomy and the continued discovery of new shrew species, as well as the expansion of the range of some shrew species^10,11^, demonstrate the high complexity of this group of animals. There is limited information available on basic parameters of shrew’s (population) biology such as population structure and dynamics. However, shrews may share similar properties with other so-called viral hyperreservoirs such as bats and rodents.^5^ Their high metabolism, torpor, fast life cycle and unknown immunological responses to viral infection may enable them to sustain and spread viral infections without developing any disease.^4,5^

Here we present an effective and robust method for deciphering the virosphere of white-toothed shrews and identified several novel viruses that are surprisingly closely related to known zoonotic and enzootic viruses of the genera *Henipavirus*, *Orthonairovirus*, *Orthohepevirus* and *Orthobornavirus*.

Viruses detected in *C. russula*, which has a North African origin, are genetically similar with other viruses detected in African shrews (ERVEV and TFAV virus for nairoviruses, and DewV and MeliV for paramyxoviruses), whereas *C. suaveolens*, which is widely distributed across Eurasia, presented viruses with close relatives detected in Asian shrews (HasV and LayV). This suggests a certain degree of co-evolution between the shrew species and their carried viruses.

In the context of increased pandemic preparedness, these viruses and their reservoirs need to be studied in more detail to assess their pathological relevance, mode of transmission, but also their potential as surrogates for vaccine development.^1,6^ Although the elusive behaviour of these synanthropic shrews makes it difficult to grasp the human-shrew interface, it does exist, as evidenced by human BoDV-1 infections. Screening of risk groups that are potentially in contact with shrews, such as agricultural workers, is recommended to assess the zoonotic potential of these viruses. This increased knowledge will help to determine the level of personal protective measures recommended when handling shrews from the wild and in captivity such as ecologists, small mammal biologists and animal keepers.^41^

Our results demonstrate the great virus diversity harboured in wildlife, not only in biodiversity hotspots ^25^, but also in industrialised countries such as in Central Europe.^2^ Finally, it is essential to decipher the virome of as many putative reservoir species as possible in order to establish novel risk models for disease emergence and preventive measures^1,46^, but the conservation of white-toothed shrews has also to be acknowledged.^3,47^ In a holistic One Health approach, these future studies should evaluate the potential influence of anthropogenic land use, biodiversity and climate change on the range of these neglected reservoir species and their potential as reservoirs for zoonotic agents.

## Contributors

Methodology, F.P., D.H. and M.B.; validation, F.P.; investigation, V.C.H., B.L., J.S.-E., D.H. and F.P.; resources, J.J., M.Ba., M.H. and M.Br.; writing—original draft preparation, V.C.H., and F.P.; writing—review and editing, V.C.H., B.L., J.S.-E., M.Ba., D.H., J.J., M.H., M.Br., R.G.U., M.B. and F.P.; visualization, F.P. and V.C.H.; conceptualization, supervision, project administration, funding acquisition, R.G.U., M.B., F.P. All authors have read and agreed to the published version of the manuscript.

## Institutional Review Board Statement

Not applicable.

## Ethical statement

Shrews were by-catches of trapping approved by State agencies (permit no: 22-2684-04-15-105/16 (GER-TH), 42502-2-1548 (UniLPZ; GER-ST), 84-02.04.2015.A279 (GER-NW), V/2/2006/10 (CZ)) and of trapping conducted by forestry authorities within their professional duties. Etruscan shrew tissue was collected according to a permit T0078/16 given to the Brecht group. The majority of small mammals originated from a Citizen Science-based project (cat prey, found dead), therefore no further permits were required.

## Informed Consent Statement

Not applicable.

## Data Availability Statement

All data are presented within the manuscript and its Supplemental materials. Viral genomes and raw read data were uploaded to GenBank using the accessions OR713845-OR713892.

## Acknowledgments

We are very grateful to Christian Imholt, Marion Saathoff, Tanja Wölk, Wolfgang Fiedler, Cornelia Triebenbacher, Karin Weber, Stefanie Zeiske-Lippert, Barbara Schmidt, Philipp Koch, Susanne Modrow, Tobias Eisenberg, Andreas Micklich, Kerstin Bauer, Kerstin Albrecht, Kirsten Pörtner, Ronny Wolf, Martin Trost, Martin Pfeffer, Nelly Scuda, Michaela Gentil and all private persons and cats, participating in our Citizen Science project for providing shrew specimens and the whole working group of Rainer Ulrich, Petra Strakova, Martina Dokulilova, and Anna R. Brück for small mammal dissection and technical support. We thank Jenny Lorke and Hanna Nitzsche for excellent technical assistance in library preparation and sequencing.

## REFERENCES

1. Dharmarajan, G. et al. The Animal Origin of Major Human Infectious Diseases: What Can Past Epidemics Teach Us About Preventing the Next Pandemic? Zoonoses 2, 1–13; 10.15212/ZOONOSES-2021-0028 (2022).

2. Jones, K. E. et al. Global trends in emerging infectious diseases. Nature 451, 990–993; 10.1038/nature06536 (2008).

3. Daszak, P., Cunningham, A. A. & Hyatt, A. D. Emerging infectious diseases of wildlife-- threats to biodiversity and human health. *Science (New York*, N.Y.) 287, 443–449; 10.1126/science.287.5452.443 (2000).

4. Luis, A. D. et al. A comparison of bats and rodents as reservoirs of zoonotic viruses: are bats special? Proceedings. Biological sciences / The Royal Society 280, 20122753; 10.1098/rspb.2012.2753 (2013).

5. Han, B. A., Schmidt, J. P., Bowden, S. E. & Drake, J. M. Rodent reservoirs of future zoonotic diseases. PNAS nexus 112, 7039–7044; 10.1073/pnas.1501598112 (2015).

6. Letko, M., Seifert, S. N., Olival, K. J., Plowright, R. K. & Munster, V. J. Bat-borne virus diversity, spillover and emergence. Nature reviews. Microbiology 18, 461–471; 10.1038/s41579-020-0394-z (2020).

7. Olival, K. J. et al. Host and viral traits predict zoonotic spillover from mammals. Nature 546, 646–650; 10.1038/nature22975 (2017).

8. 8. Wilson, D. E. & Mittermaier, R. A. (eds.). Handbook of the Mammals of the World. Volume 8 Insectivores, Sloths and Colugos (Lynx Edicions, Bellaterra (Barcelona), 2017).

9. Esselstyn, J. A. et al. Fourteen New, Endemic Species of Shrew (Genus *Crocidura*) from Sulawesi Reveal a Spectacular Island Radiation. Bulletin of the American Museum of Natural History 454, 1–108; 10.1206/0003-0090.454.1.1 (2021).

10. Bellocq, J. G. de et al. First record of the greater white-toothed shrew, Crocidura russula, in the Czech Republic. Journal of Vertebrate Biology 72, 1–9; 10.25225/jvb.23047 (2023).

11. van der Kooij, J. & Nyfors, E. Citizen science reveals the first occurrence of the greater white-toothed shrew *Crocidura russula* in Fennoscandia. Mammalia 87, 442–450; 10.1515/mammalia-2023-0042 (2023).

12. Zeller, H. G. et al. Electron microscopic and antigenic studies of uncharacterized viruses. II. Evidence suggesting the placement of viruses in the family *Bunyaviridae*. Archives of virology 108, 211–227; 10.1007/BF01310935 (1989).

13. Chastel, C. et al. Erve virus, a probable member of *Bunyaviridae* family isolated from shrews (*Crocidura russula*) in France. Acta virologica 33, 270–280 (1989).

14. Dilcher, M. et al. Genetic characterization of Erve virus, a European Nairovirus distantly related to Crimean-Congo hemorrhagic fever virus. Virus genes 45, 426–432; 10.1007/s11262-012-0796-8 (2012).

15. Ozeki, T. et al. Identification of novel orthonairoviruses from rodents and shrews in Gabon, Central Africa. Journal of General Virology 103, 1–12; 10.1099/jgv.0.001796 (2022).

16. Low, D. H. W. et al. Cencurut virus: A novel *Orthonairovirus* from Asian house shrews (*Suncus murinus*) in Singapore. *One health (Amsterdam*, Netherlands) 16, 100529; 10.1016/j.onehlt.2023.100529 (2023).

17. Hawman, D. W. & Feldmann, H. Crimean–Congo haemorrhagic fever virus. Nature reviews. Microbiology 21, 463–477; 10.1038/s41579-023-00871-9 (2023).

18. Zhang, X.-A. et al. A Zoonotic Henipavirus in Febrile Patients in China. The New England journal of medicine 387, 470–472; 10.1056/NEJMc2202705 (2022).

19. Lee, S.-H. et al. Discovery and Genetic Characterization of Novel Paramyxoviruses Related to the Genus *Henipavirus* in *Crocidura* Species in the Republic of Korea. Viruses 13, 1–16; 10.3390/v13102020 (2021).

20. Vanmechelen, B. et al. The characterization of multiple novel paramyxoviruses highlights the diverse nature of the subfamily *Orthoparamyxovirinae*. Virus evolution 8, 1–12; 10.1093/ve/veac061 (2022).

21. Chua, K. B. et al. Nipah virus: a recently emergent deadly paramyxovirus. *Science (New York*, N.Y.) 288, 1432–1435; 10.1126/science.288.5470.1432 (2000).

22. Gazal, S. et al. Nipah and Hendra Viruses: Deadly Zoonotic Paramyxoviruses with the Potential to Cause the Next Pandemic. *Pathogens (Basel*, Switzerland) 11, 1–16; 10.3390/pathogens11121419 (2022).

23. Rubbenstroth, D., Schlottau, K., Schwemmle, M., Rissland, J. & Beer, M. Human bornavirus research: Back on track! PLoS pathogens 15, e1007873; 10.1371/journal.ppat.1007873 (2019).

24. Niller, H. H. et al. Zoonotic spillover infections with Borna disease virus 1 leading to fatal human encephalitis, 1999-2019: an epidemiological investigation. Lancet Infect Dis 20, 467–477; 10.1016/S1473-3099(19)30546-8 (2020).

25. Chen, Y.-M. et al. Host traits shape virome composition and virus transmission in wild small mammals. Cell 186, 1–14; 10.1016/j.cell.2023.08.029 (2023).

26. Schlegel, M. et al. Molecular identification of small mammal species using novel *Cytochrome b* gene-derived degenerated primers. Biochemical genetics 50, 440–447; 10.1007/s10528-011-9487-8 (2012).

27. Bushnell, B. BBmap. Available at sourceforge.net/projects/bbmap/ (2023).

28. Kopylova, E., Noé, L. & Touzet, H. SortMeRNA: fast and accurate filtering of ribosomal RNAs in metatranscriptomic data. Bioinformatics (Oxford, England) 28, 3211–3217; 10.1093/bioinformatics/bts611 (2012).

29. Quast, C. et al. The SILVA ribosomal RNA gene database project: improved data processing and web-based tools. Nucleic acids research 41, D590–6; 10.1093/nar/gks1219 (2013).

30. Bushmanova, E., Antipov, D., Lapidus, A. & Prjibelski, A. D. rnaSPAdes: a de novo transcriptome assembler and its application to RNA-Seq data. GigaScience 8; 10.1093/gigascience/giz100 (2019).

31. Tamames, J. & Puente-Sánchez, F. SqueezeMeta, A Highly Portable, Fully Automatic Metagenomic Analysis Pipeline. Frontiers in microbiology 9, 3349; 10.3389/fmicb.2018.03349 (2018).

32. Martin, M. Cutadapt removes adapter sequences from high-throughput sequencing reads. EMBnet j. 17, 10; 10.14806/ej.17.1.200 (2011).

33. Minh, B. Q. et al. IQ-TREE 2: New Models and Efficient Methods for Phylogenetic Inference in the Genomic Era. Molecular biology and evolution 37, 1530–1534; 10.1093/molbev/msaa015 (2020).

34. Minh, B. Q., Nguyen, M. A. T. & Haeseler, A. von. Ultrafast approximation for phylogenetic bootstrap. Molecular biology and evolution 30, 1188–1195; 10.1093/molbev/mst024 (2013).

35. Guindon, S. et al. New algorithms and methods to estimate maximum-likelihood phylogenies: assessing the performance of PhyML 3.0. Systematic biology 59, 307–321; 10.1093/sysbio/syq010 (2010).

36. Untergasser, A. et al. Primer3--new capabilities and interfaces. Nucleic acids research 40, e115; 10.1093/nar/gks596 (2012).

37. Schlottau, K. et al. Fatal Encephalitic Borna Disease Virus 1 in Solid-Organ Transplant Recipients. The New England journal of medicine 379, 1377–1379; 10.1056/NEJMc1803115 (2018).

38. Wernike, K., Hoffmann, B., Kalthoff, D., König, P. & Beer, M. Development and validation of a triplex real-time PCR assay for the rapid detection and differentiation of wild-type and glycoprotein E-deleted vaccine strains of Bovine herpesvirus type 1. Journal of virological methods 174, 77–84; 10.1016/j.jviromet.2011.03.028 (2011).

39. Crawford, K. CDTR-Communicalbe Disease Threats Report. Weekly Report Week 38, 17–23 September 2023. Available at https://www.ecdc.europa.eu/en/publications-data/communicable-disease-threats-report-17-23-september-2023-week-38 (2023).

40. Drexler, J. F. et al. Bats host major mammalian paramyxoviruses. Nature communications 3, 1–12; 10.1038/ncomms1796 (2012).

41. Geyer, B. et al. Establishing and Maintaining an Etruscan Shrew Colony. Journal of the American Association for Laboratory Animal Science : JAALAS 61, 52–60; 10.30802/AALAS-JAALAS-21-000068 (2022).

42. Treib, J. et al. Thunderclap headache caused by Erve virus? Neurology 50, 509–511; 10.1212/wnl.50.2.509 (1998).

43. Velavan, T. P. et al. Hepatitis E: An update on One Health and clinical medicine. Liver international : official journal of the International Association for the Study of the Liver 41, 1462–1473; 10.1111/liv.14912 (2021).

44. Johne, R. et al. Novel hepatitis E virus genotype in Norway rats, Germany. Emerging Infectious Diseases 16, 1452–1455; 10.3201/eid1609.100444 (2010).

45. Negredo, A. et al. Fatal Case of Crimean-Congo Hemorrhagic Fever Caused by Reassortant Virus, Spain, 2018. Emerging Infectious Diseases 27, 1211–1215; 10.3201/eid2704.203462 (2021).

46. Zhang, Y.-Z., Chen, Y.-M., Wang, W., Qin, X.-C. & Holmes, E. C. Expanding the RNA Virosphere by Unbiased Metagenomics. Annu Rev Virol 6, 119–139; 10.1146/annurev-virology-092818-015851 (2019).

47. Sokolow, S. H. et al. Ecological interventions to prevent and manage zoonotic pathogen spillover. Philosophical Transactions of the Royal Society B: Biological Sciences 374, 1–10; 10.1098/rstb.2018.0342 (2019).

