## Supplemental for "Small in size but huge as reservoir – insights into the virome of European white-toothed shrews"

Haring *et al.* 2023

Supplemental Figures

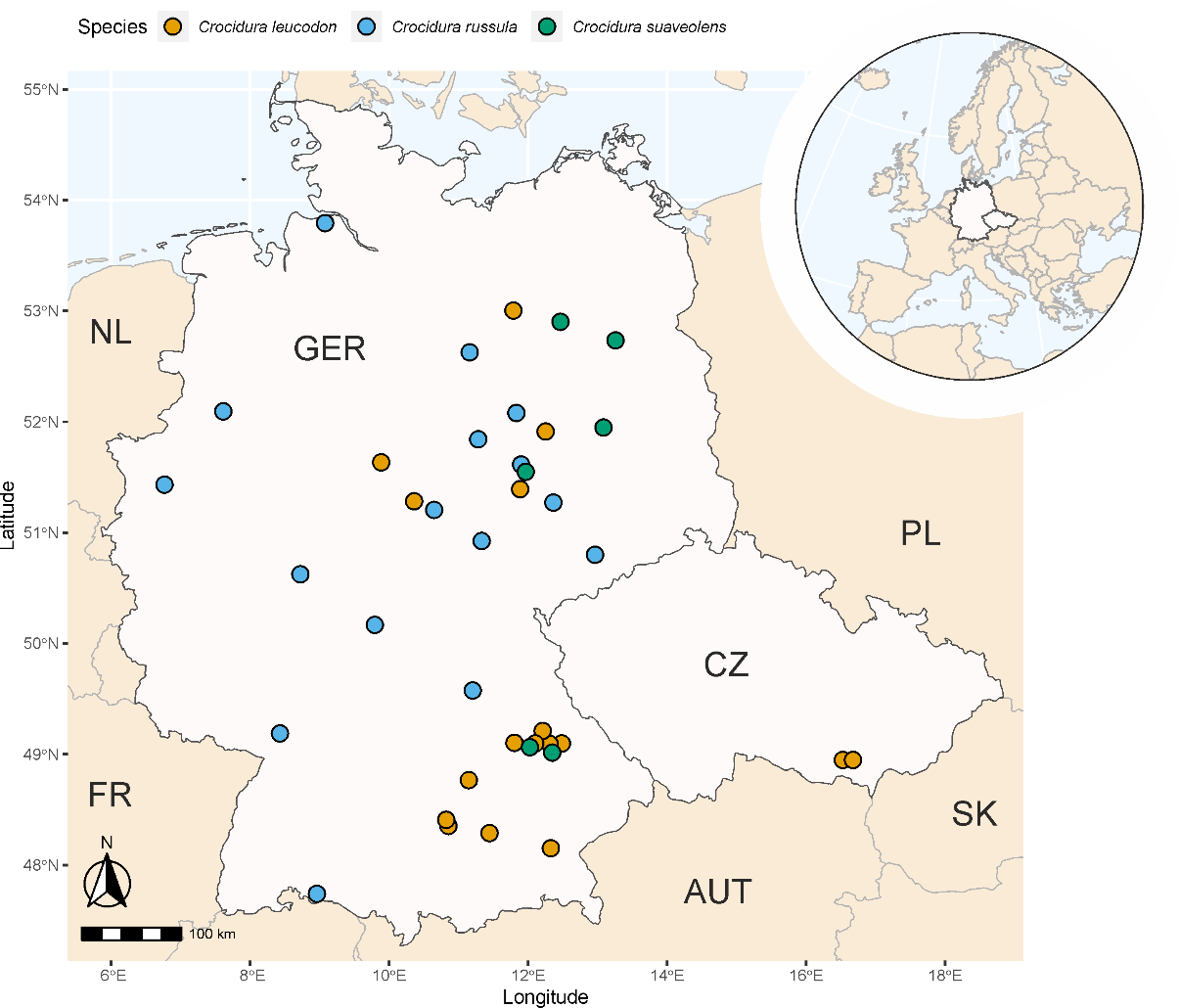

**Supplemental Figure S1**: Geographical origin of studied wild white-toothed shrews from Germany and the Czech Republic. This study included 19 bicolored white-toothed shrews (*Crocidura leucodon*), 16 greater white-toothed shrews (*Crocidura russula*), and 6 lesser white-toothed shrews (*Crocidura suaveolens*) from Germany (GER) and two *C. leucodon* collected in the Czech Republic (CZ). Two additional Etruscan shrews (*Suncus etruscus*) originated from a colony in Germany and are therefore not shown. NL: the Netherlands; FR: France; AUT: Austria; SK: Slovakia; PL: Poland

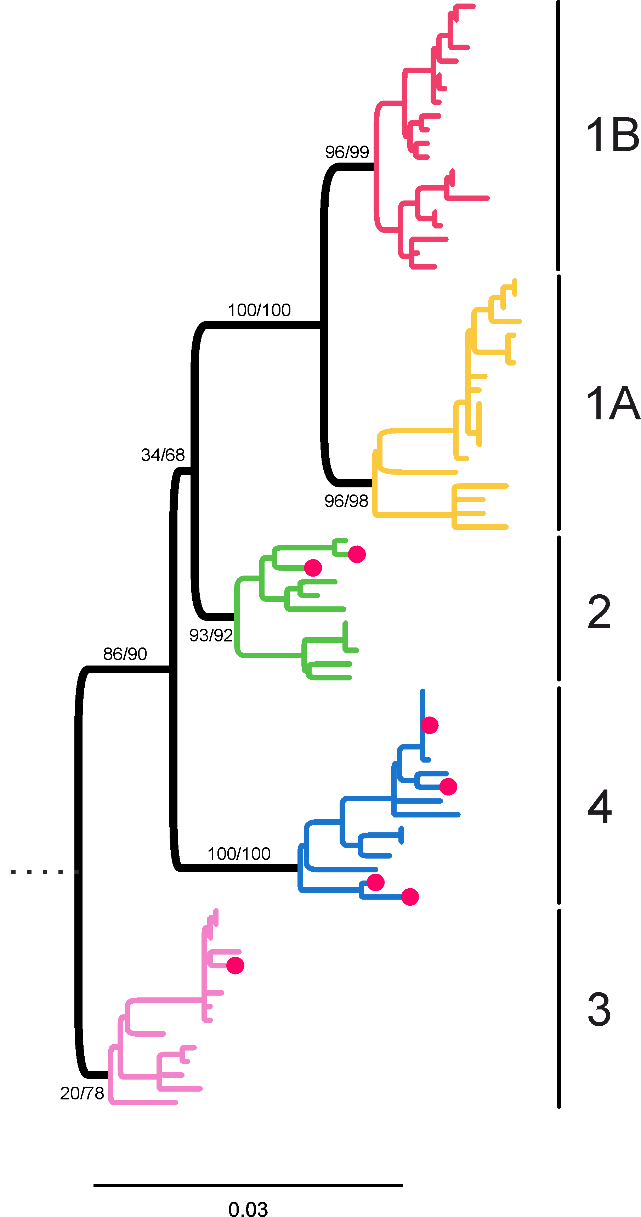

**Supplemental Figure S2**: Phylogenetic relationships of the identified BoDV-1 strains. For phylogenetic analysis, we selected 74 representative sequences of BoDV-1 together with the strains identified in this study (red dot). BoDV-2 was used as an outgroup. Only the NX/P region was selected and aligned using MUSCLE. Maximum likelihood phylogeny was inferred using IQ-TREE2 (version 2.2.2.6) with an automated model selection and 100,000 ultra-fast bootstrap and SH-aLRT replicates each. Only bootstrap values at major branches are shown. Established BoDV-1 clusters are highlighted in different colours.

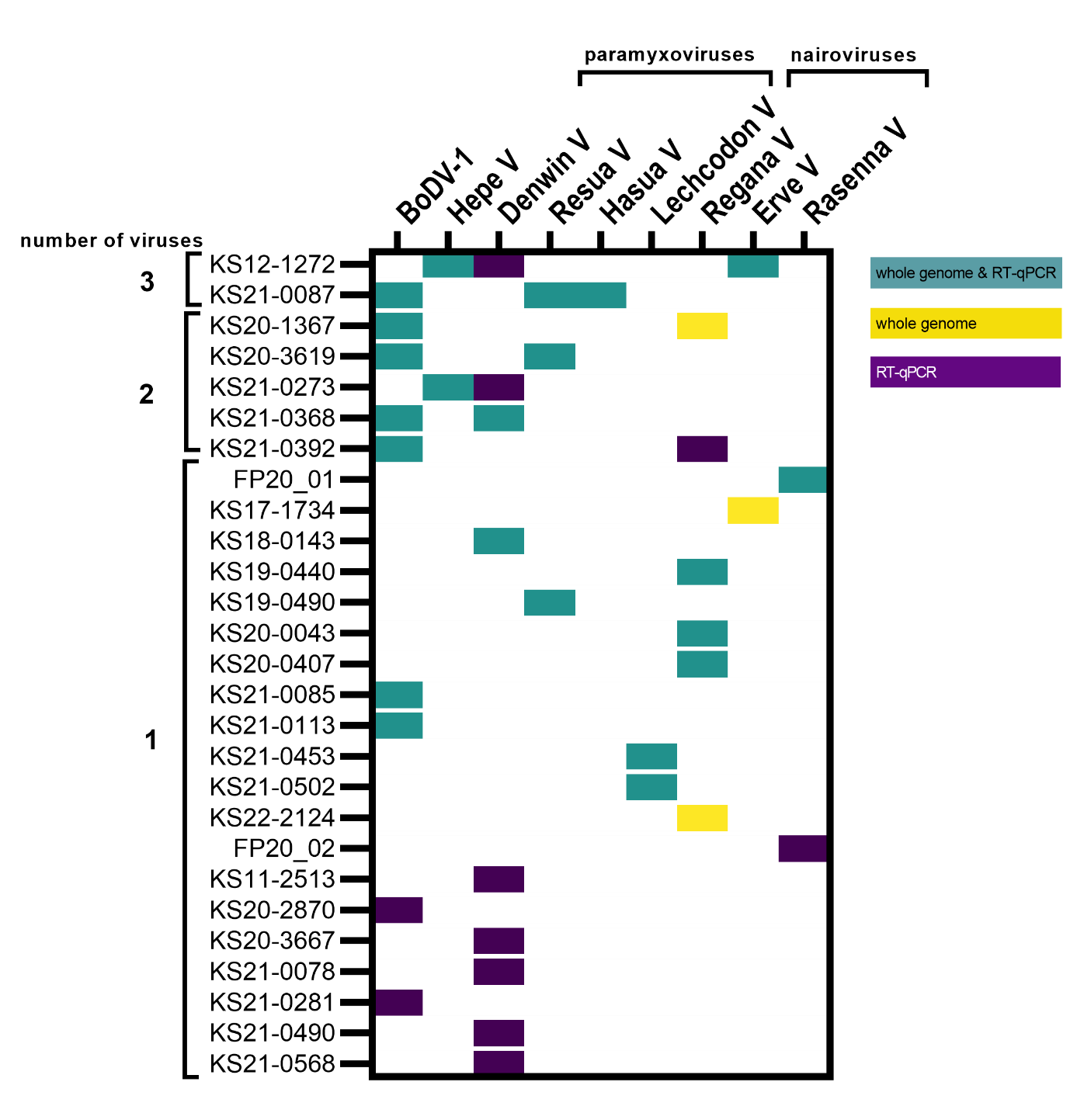

**Supplemental Figure S3**: Co-detection of up to three viruses from different families in the white-toothed shrews. Samples are sorted by the number of identified pathogens. Samples from which a viral whole genome sequence could be derived and corresponding positive RT-qPCR result are shown in turquoise, while virus identification by RT-qPCR alone is shown in purple. Two samples shown in yellow had whole genome sequences only.

Supplemental Tables

**Supplemental Table S1**: Information on the investigated white-toothed shrews, pool composition, sequencing information and detected viruses. See file “Supplemental Table S1.xlsx”.

**Supplemental Table S2**: Detailed information on the detected viruses in the white-toothed shrew samples

| **Taxonomic classification** | **Virus name** | **Tentatively named after (if novel)** | **Detected in** | **Whole genomes generated from** | **Additional RT-qPCR positive individuals** |
| --- | --- | --- | --- | --- | --- |
| *Paramyxoviridae:* *Henipavirus* | Hasua virus (HasV) | river “**Ha**vel” and the species abbreviation “*C.* ***sua****veolens*” | *C. suaveolens* | KS21-0087 | - |
|  | Resua virus (ResV) | river “**Re**gen” and the species abbreviation “*C.* ***sua****veolens*” | *C. suaveolens* | KS19-0490, KS21-0087, KS20-3619 | - |
|  | Lechcodon virus (LechV) | river “**Lech**” and the species abbreviation “*C. leu****codon****”’* | *C. leucodon* | KS21-0453, KS21-0502 | - |
|  | Denwin virus (DewV) |  | *C. russula* | KS18-0143, KS21-0368 | KS11-2513, KS12-1272, KS20-3667, KS21‑0078, KS21-0273, KS21-0490, KS21-0568 |
| *Nairoviridae: Orthonairovirus* | Rasenna virus (RASV) | named after the Etruscan civilization’s own designation “**Rasenna”** | *S. etruscus* | FP20-01 | FP20-02 |
|  | Erve virus (ERVEV) |  | *C. russula* | KS12-1272, KS17-1734 | - |
|  | Regana virus (REGV) | **“Regana”** is the Germanic word for “Regen”, the nearby river in the Bavarian region where most of the positive shrews were caught | *C. leucodon* | KS19-0440, KS20-0043, KS20-0407, KS20-1367, KS22-2124, KS21-0453 | KS21-0392 |
| *Hepeviridae: Paslahepevirus* | shrew hepatitis E virus (shrewHEV) | Phylogenetic relationship to Hepatitis E virus | *C. russula* | KS12-1272, KS21-0273 | - |
| *Bornaviridae:* *Orthobornavirus* | Borna disease virus 1 (BoDV-1, species *Orthobornavirus bornaense*) |  | *C. leucodon k*  *C. russula*  *C. suaveolens* | KS20-1367, KS21-0085, KS21-0113, KS21-0392  KS21-0368  KS21-0087, KS20-3619 | KS21-2870, KS21-0281 |

**Supplemental Table S3**: Primer and probes used for RT-qPCRs

| Name | Type | Sequence (5’→3’) |  |
| --- | --- | --- | --- |
| ReganaV-1-F | Primer | TCTCCACCTGCCTACAGAGA | This study |
| ReganaV-1-R | Primer | TGCTGCTCTTTCTTTTCAGGA | This study |
| ReganaV-1-FAM | Probe | FAM-TGCAGCAGATACTGATGGATTTCCCA-BHQ-1 | This study |
| ReganaV-2-F | Primer | CCAGAACCTAAACAGAGCATTC | This study |
| ReganaV-2-R | Primer | ACCAAATCCCACATCTGCTATA | This study |
| ReganaV-2-FAM | Probe | FAM-AATGAAGAACGCCCTTTACTTAGTGTGTG-BHQ-1 | This study |
| ERVEV-1-F | Primer | TAGAAGGTCAAGCTCATCGAAT | This study |
| ERVEV-1-R | Primer | ACTCAAGGAAAATGCCAGAATC | This study |
| ERVEV-1-FAM | Probe | FAM-ACCTCAAGTCTGATCAATAGATACACCACC-BHQ-1 | This study |
| ERVEV-2-F | Primer | AGAGGAGTTGGACAATAGGATG | This study |
| ERVEV-2-R | Primer | CTTCAAAATGCCCATTCAACAC | This study |
| ERVEV-2-FAM | Probe | FAM-CTGCTGACAAATTTTATAACTGAGGCGGTA-BHQ-1 | This study |
| RasennaV-1-F | Primer | AACGCATCATGAATGGCCACA | This study |
| RasennaV-1-R | Primer | CAAACCCAGTGGTAAGCAGCA | This study |
| RasennaV-1-FAM | Probe | FAM-CCACCTTGGGGAGATGTGGATAAGCA-BHQ-1 | This study |
| LechcodonV-F | Primer | TACATCCAACTGAACATGAACT | This study |
| LechcodonV-R | Primer | ATCATTAATCTCCCTTCAAGCA | This study |
| LechcodonV-FAM | Probe | FAM-GTGTCGTTGATTGGAATAGTCGAAAGATCT-BHQ-1 | This study |
| DewV-F | Primer | CAGAAACAATAATCAGTACACATTTC | This study |
| DewV-R | Primer | CACCTTGATATAGATTTTAGTCCC | This study |
| DewV-FAM | Probe | FAM-TTCCAAAAGGATTTACTATGATGGGATGGT-BHQ-1 | This study |
| HasuaV-F | Primer | TCAGATAAACATGAACCTTGTC | This study |
| HasuaV-R | Primer | TCACAATACATGAGAACAAGTT | This study |
| HasuaV-FAM | Probe | FAM-GGAACAAGAAACAAATTTATCATTTAACACCA-BHQ-1 | This study |
| ResuaV-F | Primer | TCAATCAAAATCTTTGCACCAT | This study |
| ResuaV-R | Primer | CCTTCAACAACATCACAATACA | This study |
| ResuaV-FAM | Probe | FAM-AGAGATTTACCATTTAACTCCTGAACTTGT-BHQ-1 | This study |
| shrewHEV-F | Primer | CAGACGCGGTGGTTCAAAC | This study |
| shrewHEV-R | Primer | GTGGAACCAAGGGCAGCT | This study |
| shrewHEV-FAM | Probe | FAM-CCAGCCAGAGTCATTTCCACTAACAACCC-BHQ-1 | This study |
| BoDV-1-1288-F | Primer | TAGTYAGGAGGCTCAATGGCA | ^1^ |
| BoDV-1-1449-R | Primer | GTCCYTCAGGAGCTGGTC | ^1^ |
| BoDV-1-1346-FAM | Probe | FAM-AAGAAGATCCCCAGACACTACGACG-BHQ1 | ^1^ |
| ACT-1030-F | Primer | AGCGCAAGTACTCCGTGTG | ^2^ |
| ACT-1135-R | Primer | CGGACTCATCGTACTCCTGCTT | ^2^ |
| ACT-1081-HEX | Probe | HEX-TCGCTGTCCACCTTCCAGCAGATGT-BHQ1 | ^2^ |

Supplemental References

1.	Schlottau, K. *et al.* Fatal Encephalitic Borna Disease Virus 1 in Solid-Organ Transplant Recipients. *The New England journal of medicine* **379,** 1377–1379; 10.1056/NEJMc1803115 (2018).

2.	Wernike, K., Hoffmann, B., Kalthoff, D., König, P. & Beer, M. Development and validation of a triplex real-time PCR assay for the rapid detection and differentiation of wild-type and glycoprotein E-deleted vaccine strains of Bovine herpesvirus type 1. *Journal of virological methods* **174,** 77–84; 10.1016/j.jviromet.2011.03.028 (2011).
